# GatorPrism: Prototype-Conditioned Routing across Coalition Graph Experts for Spatial Multi-Omics Integration

**DOI:** 10.64898/2026.07.21.739865

**Authors:** Zhenhao Zhang, Yiyao Zhang, Xiangtao Li, Jiang Bian, Jun Shen, Yuxi Liu

## Abstract

Spatial multi-omics integration requires balancing cross-omics consensus with modality-specific signals whose importance varies across tissue locations. We present GatorPrism, a self-supervised coalition graph mixture-of-experts framework that explicitly separates cross-omics consensus from modality-specific structure and adaptively integrates their contributions across tissue locations. A joint expert encodes an intersection-based consensus graph, while modality-private experts capture mixed spatial–molecular structures; a prototype-conditioned router assigns spot-specific coalition weights. GatorPrism is trained end-to-end to preserve shared and modality-specific neighborhoods, align co-registered modalities, maintain spatial coherence, and prevent routing collapse. Across eight spatial multi-omics datasets, GatorPrism achieved strong performance across nine clustering metrics against nine competing methods. In human tonsil, inferred domains were supported by concordant RNA and ADT markers and distinct functional programs. In embryonic mouse brain, routing profiles revealed anatomically localized shared, RNA-private, and ATAC-private states supported by transcriptomic, chromatin-accessibility, and motif evidence. These results establish GatorPrism as an accurate and interpretable framework that reveals how shared and modality-specific molecular signals organize tissue structure. Source code is available at https://github.com/Gator-Group/GatorPrism.

## 1 Introduction

Spatially resolved multi-omics technologies enable the joint measurement of multiple molecular layers while preserving their tissue coordinates [8, 10, 17]. By profiling transcriptomic, proteomic, epigenomic, and other molecular signals within the same spatial context, these technologies provide a multidimensional characterization of cellular identity, regulatory states, and tissue function [12]. These multimodal profiles are particularly valuable for identifying spatial domains, which are coherent tissue regions characterized by distinct molecular programs, cellular compositions, or microenvironmental states. Effective spatial domain identification can advance the study of developmental organization [24], immune architecture [25], tumor heterogeneity [22, 29], cellular niche organization [3], and disease-associated tissue remodeling [32]. However, fully leveraging these data requires computational methods that can integrate heterogeneous molecular measurements while preserving both spatial organization and modality-specific biological signals.

Spatial multi-omics integration remains challenging because omics modalities differ substantially in dimensionality, sparsity, noise, and statistical distribution, while each captures a distinct layer of cellular state and regulation [1]. Spatial proximity and molecular similarity are also not equivalent [7, 20, 33]: neighboring spots may belong to different tissue compartments, whereas spatially distant spots may share similar molecular profiles. Moreover, molecular neighborhoods can vary across modalities [10, 13, 14], requiring integration methods to preserve spatial coherence while capturing both shared cross-omics structure and modality-specific molecular organization.

To address these challenges, existing methods learn unified representations by joining, aligning, or modeling interactions among modality-specific embeddings [6, 16, 27]; adaptively weighting modality-specific representations [19, 35]; learning modality-specific graph structures before fusion [9]; or deriving spatial consensus representations through statistical or ensemble modeling [4, 15]. SpaMV instead explicitly decomposes spatial multi-omics data into cross-omics shared and omics-specific private representations [18].

Despite these advances, existing methods do not determine whether an integrated representation should rely primarily on cross-omics consensus or modality-specific evidence at each spatial location. SpatialGlue [19] and COSMOS [35] learn spot-specific weights over modality representations, but do not explicitly distinguish molecular relationships jointly supported across modalities from those unique to an individual modality. SpaMV decomposes shared and private representations, but does not adaptively estimate their relative contributions across spatial locations [18]. Accordingly, current methods do not explicitly characterize whether a tissue region is organized by concordant multi-omic signals or mainly by one modality. The central problem is therefore to learn an integrated spatial representation whose reliance on cross-omics consensus and modality-specific evidence varies across locations.

Addressing this problem requires cross-omics consensus and modality-specific information to be distinguished prior to adaptive fusion, beginning at the graph-construction stage. Existing methods typically encode each modality using its own feature or mixed neighborhood graph together with spatial structure [9, 18, 19], but do not jointly construct semantically distinct consensus and modality-specific graphs. As a result, their fusion mechanisms cannot explicitly attribute integrated representations to relationships shared across modalities or to modality-specific molecular structure. These limitations motivate the separate representation of these information sources and their spot-specific integration, providing a principled basis for interpreting spatial variation in their relative contributions.

Here, we present GatorPrism, an interpretable self-supervised coalition graph mixture-of-experts framework for spatial multiomics integration (Fig. 1). GatorPrism distinguishes cross-omics consensus from modality-specific molecular structure at the graph-construction stage. A joint expert encodes concatenated multiomics features over an intersection-based consensus graph, whereas modality-private experts encode individual modalities over mixed graphs combining spatial proximity with modality-specific molecular neighborhoods. A prototype-conditioned router uses the latent context of each spatial location to learn spot-specific coalition weights and adaptively fuse the expert representations. GatorPrism is trained end-to-end using modality-aware reconstruction, consensus and modality-specific neighborhood contrast, cross-modal alignment, spatial coherence, and expert load balancing. The fused representation supports spatial domain identification, while domain-level aggregation of routing weights reveals localized reliance on shared and modality-specific information.

**Figure 1:**
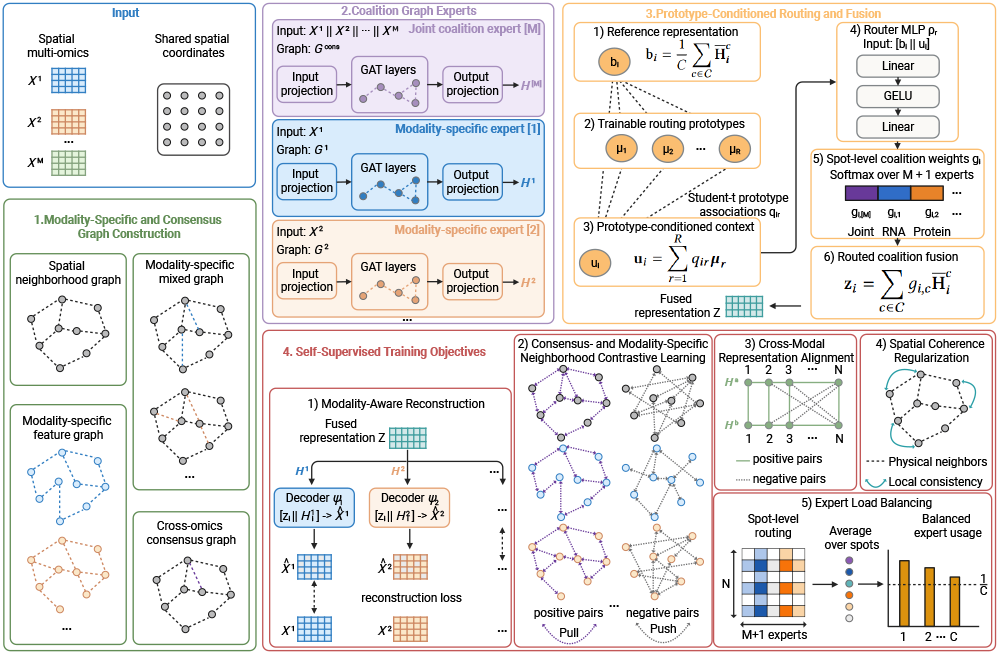
Overview of the GatorPrism framework.

Our main contributions are summarized as follows:

- We formulate spatial multi-omics integration as a spatially adaptive representation learning problem in which the relative contributions of cross-omics consensus and modality-specific signals vary across tissue locations.
- We introduce a coalition graph mixture-of-experts architecture with semantically distinct joint and modality-private experts, together with prototype-conditioned routing for spot-specific expert integration.
- We develop a unified self-supervised objective that preserves consensus and modality-specific neighborhood structures, promotes cross-modal alignment and spatial coherence, and prevents routing collapse.
- We benchmark GatorPrism against nine competing methods across eight spatial multi-omics datasets, demonstrating robust performance and revealing anatomically localized shared-, RNA-private-, and ATAC-private routing states supported by coherent molecular and regulatory evidence.

## 2 Notation and Problem Formulation

We consider unsupervised spatial domain identification from a spatially co-registered multi-omics tissue section with *N* spatial units and *M* ≥ 2 molecular modalities. We use [*N*] = {1, …, *N*} and [*M*] = {1, …, *M*} to index the spatial units and modalities, respectively. For each modality *m* = 1, …, *M*, let 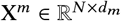 denote its preprocessed feature matrix, whose *i*-th row, 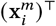, denotes the feature vector of spatial unit *i* in modality *m*. Let S ∈ ℝ^*N*×2^ denote the corresponding spatial coordinate matrix. Accordingly, the input can be written as

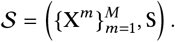

Because the modalities are spatially co-registered, row *i* of every X^*m*^ and row *i* of S correspond to the same spatial unit.

Given *S* and a prespecified number of spatial domains *K*, the objective is to partition the *N* spatial units into *K* spatially coherent domains without using spatial domain annotations. GatorPrism learns a fused representation Z ∈ ℝ^*N* ×*d*^, whose *i*-th row, 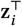, summarizes the multi-omic and spatial information associated with spatial unit *i*. The final domain assignments are obtained as

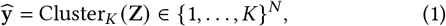

where Cluster_*K*_ denotes a *K*-component clustering procedure. In GatorPrism, this procedure is implemented using the clustering method provided by `mclust` [21].

## 3 Method

### 3.1 Modality-Specific and Consensus Graph Construction

Spatial multi-omics data encode physical proximity and modality-specific molecular similarity. Because molecular neighborhoods may differ across modalities, GatorPrism constructs modality-specific mixed graphs and a cross-omics consensus graph rather than a single fused graph.

All graphs share the node set *V* = {*v*_1_, …, *v*_*N*_}, where each node denotes a spatial unit. Given a node representation A ∈ ℝ^*N* ×*p*^, let *N*_*κ*_ (*i*; A) denote the indices of the *κ* nearest neighbors of node *i* under Euclidean distance, excluding *i*. Accordingly, we define the symmetrized *κ*-nearest-neighbor edge set:

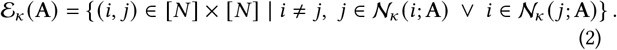

#### Spatial neighborhood graph

We construct a spatial graph *G*_sp_ =(*V, E*_sp_) from the spot coordinates S ∈ ℝ^*N* ×2^, with

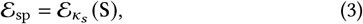

where *k*_*s*_ is the spatial neighborhood size.

#### Modality-specific feature and mixed graphs

To capture modality-dependent molecular structure, we construct the feature edge set for each modality *m*:

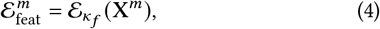

where *κ*_*f*_ is the molecular neighborhood size. These edges are then combined with the shared spatial edges to form the modality-specific mixed graph:

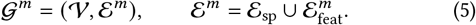

#### Cross-omics consensus graph

To represent molecular-neighborhood agreement across modalities, we retain only feature edges shared by all modality-specific feature graphs:

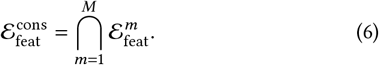

The consensus graph is defined:

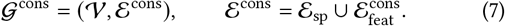

Thus, the consensus graph retains all spatial edges and only feature edges supported by every modality.

### 3.2 Coalition Graph Experts and Prototype-Conditioned Routing

Given the graph family in Section 3.1, GatorPrism assigns each omics coalition to a graph expert and integrates the resulting representations using a prototype-conditioned router. This yields a fused representation and spot-specific coalition weights.

*Coalition Graph Experts*. GatorPrism defines the coalition set:

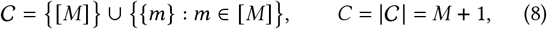

comprising one joint expert for the grand coalition [*M*] and one modality-specific expert for each singleton {*m*}. Intermediate modality subsets are not instantiated.

C The input associated with coalition *c* ∈ C is

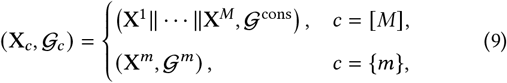

where ∥ denotes feature concatenation. Each coalition is encoded by a separate graph attention encoder_*fc*_ [28]:

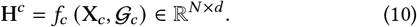

All experts therefore generate representations in a common latent space.

#### Prototype-Conditioned Routing and Fusion

Rather than using a globally fixed mixture, GatorPrism learns spot-specific coalition weights. To place the outputs of different experts on comparable scales, we normalize each expert representation:

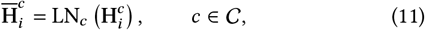

where LN_*c*_ denotes an expert-specific layer-normalization operation. We then compute the coalition-averaged routing reference:

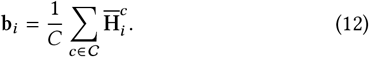

Let 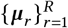 denote *R* trainable routing prototypes. Using a Student-*t* kernel [30], the soft association between spot *i* and prototype *r* is computed as:

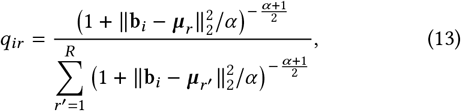

where α *>* 0 is the degrees-of-freedom parameter. The prototype-conditioned routing context is then computed as

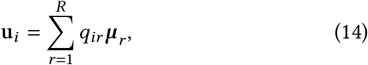

where *q*_*ir*_ is an internal routing association rather than an assignment to a spatial domain. GatorPrism does not use target sharpening or a clustering KL objective.

A two-layer multilayer perceptron *ρ*_r_ generates the spot-specific coalition weights:

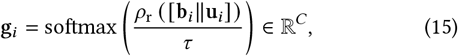

where *g*_*i,c*_ ≥ 0, ∑_*c*_ *g*_*i,c*_ = 1, and τ *>* 0 controls the sharpness of the routing distribution.

The fused representation is obtained as a convex combination of the normalized expert representations:

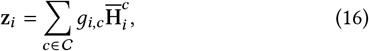

where *g*_*i,c*_ directly specifies the relative contribution of expert *c* to the fused representation at spot *i*.

### 3.3 Self-Supervised Training Objectives

GatorPrism jointly optimizes coalition experts, routing prototypes, prototype-conditioned router, fusion module, and modality-specific decoders without spatial domain annotations using self-supervised objectives.

#### Modality-Aware Reconstruction

For each modality *m*, a decoder ψ_*m*_ reconstructs its processed feature vector from the fused and modality-specific representations:

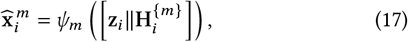

where 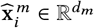 reconstructs 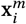. This design encourages the two latent spaces to retain complementary information. The reconstruction loss then is:

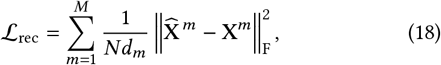

where normalization by *Nd*_*m*_ limits the influence of differences in the dimensionality of the modality.

#### Consensus- and Modality-Specific Neighborhood Contrastive Learning

GatorPrism applies contrastive objectives [23, 34] to the fused representation Z and modality-specific representations 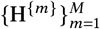 .

We normalize the representations:

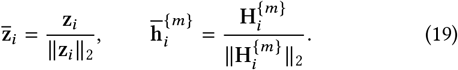

The temperature-scaled cosine similarities are:

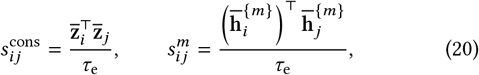

where τ_e_ *>* 0 is the edge-contrastive temperature.

For the fused representation, positive pairs are defined by the consensus feature edges, with spatial edges used when the consensus set is empty:

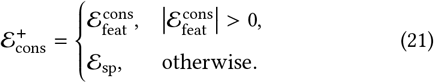

An equal number of distinct pairs of non-edge spots are uniformly sampled without replacement from the complement of 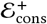 to form the negative set 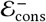. The consensus contrastive loss is:

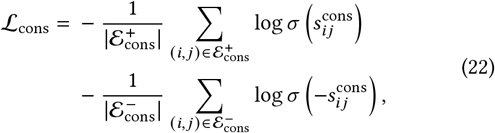

where τ (·) is the logistic sigmoid.

For modality *m*, positive pairs are given by 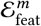, and an equal number of distinct non-edge spot pairs are uniformly sampled without replacement from the complement of 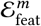 to form 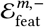.

The modality-specific neighborhood loss is

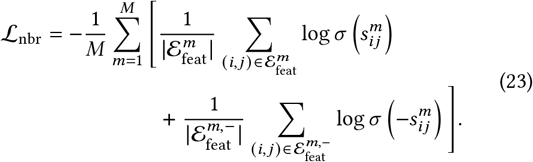

Overall, *L*_cons_ and *L*_nbr_ preserve cross-modal consensus structure in the fused space and modality-specific molecular geometry in the private expert spaces, respectively.

#### Cross-Modal Representation Alignment

For each modality pair τ *<* τ, we apply a symmetric cross-modal NT-Xent objective:

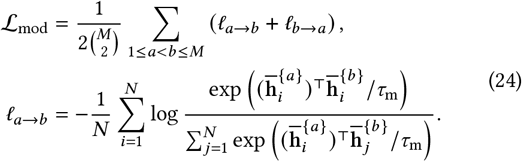

where τ_m_ *>* 0 is the cross-modal contrastive temperature, which controls the concentration of the cross-modal similarity distribution. The reverse term is obtained by exchanging *a* and *b*. Co-registered representations form positive pairs, whereas the remaining spatial units serve as cross-modal in-batch negatives [5].

#### Spatial Coherence Regularization

We encourage adjacent spatial units to have similar fused representations using graph Dirichlet energy [2]:

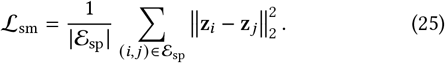

#### Expert Load Balancing

To prevent routing collapse, let the average weight assigned to expert *c* be

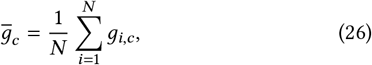

and define

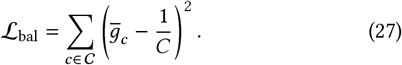

This term balances aggregate expert usage without enforcing uniform routing at each spatial unit.

#### Joint Optimization Objective

The complete objective is:

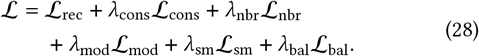

The non-negative coefficients control the auxiliary objectives, while *L*_rec_ is assigned unit weight.

### 3.4 Spatial Domain Identification and Coalition Profiling

#### Spatial Domain Identification

After training, a *K*-component model-based clustering procedure implemented in `mclust` [21] is applied to Z. The resulting component assignments define the spatial domain labels.

#### Domain-Level Coalition Profiling

Let 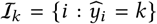 denote the spatial units assigned to domain τ. Its coalition profile is

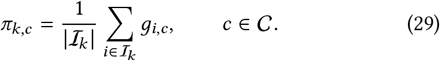

The normalized vector π_*k*_ = (π_*k,c*_)_,*c c*∈*C*_ summarizes the domain’s average routing allocation across joint and modality-specific experts.

## 4 Experiments

### 4.1 Benchmark Datasets, Baselines, and Evaluation Metrics

We evaluated GatorPrism on eight spatial multi-omics datasets: six public datasets and two simulated datasets spanning RNA+ADT and RNA+ATAC settings (see Appendix A.1). The public datasets include human lymph node (GSE263617), human tonsil (Zenodo record 18946723), P22 mouse brain from the UCSC Cell Browser, and embryonic mouse brain MISAR-seq data (OEP003285). The simulated datasets were generated following the nonnegative spatial factorization framework [26]. We compared GatorPrism with nine representative methods: CellCharter [27], SpatialGlue [19], MISO [6], COSMOS [35], PRAGA [9], SMOPCA [4], SMODEL [15],

PRESENT [16], and SpaMV [18]. Baseline descriptions are provided in Appendix B.1. Performance was evaluated using ARI, AMI, NMI, FMI, homogeneity, purity, completeness, Jaccard index, and Dice index, with higher values indicating stronger agreement with reference annotations. **Dataset links, simulation details, source code, and enlarged figures are available on GitHub, and ablation studies, hyperparameter sensitivity analysis, and other implementation details are provided in the appendix**.

### 4.2 Experimental Results

#### 4.2.1 GatorPrism achieves superior performance across spatial multiomics benchmarks

We compared GatorPrism with existing spatial multi-omics integration methods across eight benchmark datasets (Tables 1 and 2). GatorPrism achieved the highest ARI on every dataset and strong performance across the remaining clustering metrics. On the real RNA+ADT datasets, its ARI ranged from 0.3223 to 0.3917. The improvement was particularly clear on mouse brain, where GatorPrism achieved an ARI of 0.5719 compared with 0.3840 for the strongest baseline, while it reached ARIs of 0.9976 and 0.9555 on the simulated RNA+ADT and RNA+ATAC datasets, respectively. These results demonstrate that GatorPrism provides robust spatial domain identification across tissue types, species, and modality pairs.

**Table 1:** Performance comparison on RNA+ADT spatial multi-omics benchmark datasets. Values are reported as mean (standard deviation). Best and second-best results are shown in bold and underlined, respectively.

| Methods | ARI | AMI | NMI | FMI | Homo | Purity | Completeness | Jaccard | Dice |
| --- | --- | --- | --- | --- | --- | --- | --- | --- | --- |
| *Human Lymph Node-A1* |  |  |  |  |  |  |  |  |  |
| CellCharter | 0.1857(0.0135) | 0.3387(0.0114) | 0.3430(0.0113) | 0.3391(0.0141) | 0.3822(0.0108) | 0.5556(0.0182) | 0.3112(0.0124) | 0.1982(0.0115) | 0.3307(0.0159) |
| SpatialGlue | 0.2473(0.0157) | 0.3717(0.0106) | 0.3758(0.0106) | 0.3940(0.0161) | 0.4167(0.0097) | 0.6274(0.0209) | 0.3424(0.0123) | 0.2400(0.0142) | 0.3869(0.0187) |
| MISO | 0.1412(0.0459) | 0.2706(0.0469) | 0.2764(0.0463) | **0.4729(0.0267)** | 0.2161(0.0529) | 0.4923(0.0393) | 0.3985(0.0430) | 0.2682(0.0166) | 0.4227(0.0206) |
| COSMOS | 0.1608(0.0257) | 0.2986(0.0231) | 0.3030(0.0229) | 0.3078(0.0247) | 0.3492(0.0256) | 0.5448(0.0161) | 0.2676(0.0210) | 0.1715(0.0174) | 0.2924(0.0255) |
| PRAGA | 0.2655(0.0135) | 0.3693(0.0150) | 0.3734(0.0149) | 0.4097(0.0120) | 0.4133(0.0208) | 0.6349(0.0219) | 0.3407(0.0118) | 0.2521(0.0108) | 0.4026(0.0137) |
| SMOPCA | 0.1745(0.0157) | 0.2734(0.0178) | 0.2787(0.0177) | 0.3978(0.0326) | 0.2656(0.0177) | 0.5290(0.0170) | 0.2946(0.0271) | 0.2432(0.0192) | 0.3909(0.0251) |
| SMODEL | 0.2651(0.0024) | 0.3973(0.0023) | 0.4013(0.0023) | 0.4143(0.0022) | 0.4329(0.0023) | 0.6590(0.0031) | 0.3739(0.0028) | 0.2584(0.0019) | 0.4107(0.0025) |
| PRESENT | 0.2357(0.0179) | 0.3756(0.0139) | 0.3796(0.0139) | 0.3815(0.0159) | 0.4256(0.0149) | 0.6273(0.0257) | 0.3426(0.0136) | 0.2290(0.0128) | 0.3724(0.0169) |
| SpaMV | 0.2650(0.0169) | 0.3924(0.0113) | 0.3965(0.0113) | 0.4128(0.0184) | 0.4317(0.0082) | 0.6325(0.0179) | 0.3667(0.0154) | 0.2564(0.0166) | 0.4079(0.0209) |
| GatorPrism | **0.3223(0.0242)** | **0.4396(0.0186)** | **0.4429(0.0185)** | 0.4564(0.0297) | **0.4859(0.0198)** | **0.6716(0.0279)** | **0.4069(0.0209)** | **0.2893(0.0262)** | **0.4488(0.0342)** |
| *Human Tonsil-S1* |  |  |  |  |  |  |  |  |  |
| CellCharter | 0.1241(0.0246) | 0.2180(0.0356) | 0.2187(0.0356) | 0.4434(0.0200) | 0.2379(0.0417) | 0.6398(0.0251) | 0.2026(0.0316) | 0.2805(0.0176) | 0.4379(0.0213) |
| SpatialGlue | 0.1762(0.0509) | 0.2557(0.0357) | 0.2564(0.0357) | 0.4696(0.0440) | 0.2833(0.0335) | 0.6672(0.0214) | 0.2343(0.0370) | 0.3015(0.0404) | 0.4620(0.0468) |
| MISO | 0.1904(0.1000) | 0.2286(0.0570) | 0.2294(0.0569) | 0.5228(0.0851) | 0.2248(0.0467) | 0.6712(0.0424) | 0.2363(0.0739) | 0.3560(0.0815) | 0.5206(0.0834) |
| COSMOS | 0.0510(0.0311) | 0.1268(0.0410) | 0.1276(0.0410) | 0.4083(0.0374) | 0.1380(0.0476) | 0.6075(0.0198) | 0.1193(0.0368) | 0.2538(0.0320) | 0.4040(0.0394) |
| PRAGA | 0.1875(0.0481) | 0.2774(0.0222) | 0.2781(0.0222) | 0.4875(0.0416) | 0.3019(0.0246) | 0.6551(0.0113) | 0.2581(0.0229) | 0.3186(0.0388) | 0.4821(0.0431) |
| SMOPCA | 0.0124(0.0461) | 0.1054(0.0387) | 0.1064(0.0386) | 0.4973(0.0385) | 0.0945(0.0385) | 0.6128(0.0034) | 0.1238(0.0385) | 0.3254(0.0288) | 0.4903(0.0324) |
| SMODEL | 0.1215(0.0025) | 0.2436(0.0012) | 0.2442(0.0012) | 0.4367(0.0031) | 0.2693(0.0012) | 0.6573(0.0014) | 0.2234(0.0016) | 0.2743(0.0028) | 0.4305(0.0034) |
| PRESENT | 0.1264(0.0631) | 0.2417(0.0431) | 0.2425(0.0430) | 0.4792(0.0301) | 0.2518(0.0550) | 0.6533(0.0383) | 0.2350(0.0334) | 0.3130(0.0276) | 0.4762(0.0322) |
| SpaMV | 0.2514(0.0712) | 0.3009(0.0311) | 0.3016(0.0311) | 0.5425(0.0577) | 0.3184(0.0293) | 0.6896(0.0237) | 0.2869(0.0340) | 0.3718(0.0562) | 0.5399(0.0598) |
| GatorPrism | **0.3917(0.0762)** | **0.3664(0.0458)** | **0.3674(0.0458)** | **0.6369(0.0593)** | **0.4002(0.0512)** | **0.7462(0.0356)** | **0.3395(0.0434)** | **0.4664(0.0559)** | **0.6361(0.0627)** |
| *Human Tonsil-S2* |  |  |  |  |  |  |  |  |  |
| CellCharter | 0.1599(0.0143) | 0.2356(0.0121) | 0.2366(0.0121) | 0.4329(0.0079) | 0.2775(0.0177) | 0.6657(0.0305) | 0.2063(0.0095) | 0.2670(0.0086) | 0.4214(0.0107) |
| SpatialGlue | 0.1663(0.0248) | 0.2498(0.0259) | 0.2508(0.0259) | 0.4345(0.0217) | 0.2957(0.0304) | 0.6777(0.0166) | 0.2178(0.0229) | 0.2677(0.0190) | 0.4220(0.0238) |
| MISO | 0.2261(0.0496) | 0.2733(0.0299) | 0.2743(0.0299) | 0.5108(0.0536) | 0.2956(0.0197) | 0.7186(0.0116) | 0.2575(0.0443) | 0.3411(0.0518) | 0.5068(0.0531) |
| COSMOS | 0.0643(0.0308) | 0.1399(0.0373) | 0.1410(0.0373) | 0.3560(0.0213) | 0.1683(0.0454) | 0.5937(0.0193) | 0.1213(0.0316) | 0.2079(0.0157) | 0.3440(0.0213) |
| PRAGA | 0.1941(0.0379) | 0.2636(0.0108) | 0.2646(0.0108) | 0.4607(0.0379) | 0.3062(0.0138) | 0.6830(0.0121) | 0.2332(0.0122) | 0.2915(0.0366) | 0.4503(0.0422) |
| SMOPCA | 0.1328(0.0506) | 0.2271(0.0237) | 0.2282(0.0237) | 0.5041(0.0365) | 0.2249(0.0277) | 0.6648(0.0317) | 0.2327(0.0259) | 0.3359(0.0320) | 0.5022(0.0341) |
| SMODEL | 0.1403(0.0008) | 0.2145(0.0009) | 0.2155(0.0009) | 0.4293(0.0009) | 0.2469(0.0008) | 0.6703(0.0010) | 0.1912(0.0010) | 0.2674(0.0008) | 0.4219(0.0010) |
| PRESENT | 0.1753(0.0158) | 0.2560(0.0081) | 0.2569(0.0081) | 0.4648(0.0194) | 0.2886(0.0085) | 0.6807(0.0257) | 0.2317(0.0096) | 0.2989(0.0183) | 0.4599(0.0217) |
| SpaMV | 0.1936(0.0270) | 0.2695(0.0211) | 0.2705(0.0211) | 0.4644(0.0265) | 0.3106(0.0308) | 0.6890(0.0226) | 0.2399(0.0170) | 0.2952(0.0258) | 0.4553(0.0305) |
| GatorPrism | **0.3706(0.0849)** | **0.3413(0.0646)** | **0.3422(0.0645)** | **0.6112(0.0753)** | **0.3720(0.0662)** | **0.7466(0.0403)** | **0.3168(0.0625)** | **0.4392(0.0673)** | **0.6104(0.0799)** |
| *Simulation Dataset (RNA+ADT)* |  |  |  |  |  |  |  |  |  |
| CellCharter | 0.9732(0.0650) | 0.9767(0.0442) | 0.9768(0.0440) | 0.9823(0.0432) | 0.9788(0.0415) | 0.9863(0.0364) | 0.9747(0.0464) | 0.9680(0.0759) | 0.9822(0.0435) |
| SpatialGlue | 0.9189(0.0764) | 0.9170(0.0453) | 0.9174(0.0451) | 0.9489(0.0436) | 0.9211(0.0651) | 0.9696(0.0331) | 0.9149(0.0250) | 0.9041(0.0752) | 0.9480(0.0456) |
| MISO | 0.5927(0.1327) | 0.7119(0.1005) | 0.7132(0.1000) | 0.7234(0.0936) | 0.7440(0.1029) | 0.8434(0.0505) | 0.6851(0.0984) | 0.5674(0.1057) | 0.7183(0.0937) |
| COSMOS | 0.3795(0.0350) | 0.5875(0.0164) | 0.5892(0.0163) | 0.5597(0.0273) | 0.6483(0.0155) | 0.7428(0.0194) | 0.5401(0.0167) | 0.3727(0.0269) | 0.5425(0.0280) |
| PRAGA | 0.6276(0.1171) | 0.7591(0.0507) | 0.7602(0.0505) | 0.7572(0.0827) | 0.7582(0.0663) | 0.8821(0.0302) | 0.7652(0.0562) | 0.6127(0.1137) | 0.7545(0.0838) |
| SMOPCA | 0.9196(0.0000) | 0.8970(0.0000) | 0.8975(0.0000) | 0.9474(0.0000) | 0.9007(0.0000) | 0.9715(0.0000) | 0.8943(0.0000) | 0.9001(0.0000) | 0.9474(0.0000) |
| SMODEL | 0.9564(0.1207) | 0.9683(0.0710) | 0.9685(0.0707) | 0.9712(0.0799) | 0.9687(0.0665) | 0.9882(0.0314) | 0.9683(0.0748) | 0.9528(0.1270) | 0.9712(0.0801) |
| PRESENT | 0.2745(0.0383) | 0.5362(0.0292) | 0.5383(0.0290) | 0.4986(0.0198) | 0.5667(0.0399) | 0.7485(0.0227) | 0.5130(0.0223) | 0.3275(0.0156) | 0.4932(0.0175) |
| SpaMV | 0.9241(0.0734) | 0.9274(0.0665) | 0.9278(0.0661) | 0.9531(0.0444) | 0.9128(0.0803) | 0.9728(0.0264) | 0.9438(0.0506) | 0.9119(0.0818) | 0.9522(0.0456) |
| GatorPrism | **0.9976(0.0009)** | **0.9966(0.0014)** | **0.9966(0.0014)** | **0.9981(0.0007)** | **0.9967(0.0015)** | **0.9992(0.0003)** | **0.9965(0.0013)** | **0.9962(0.0014)** | **0.9981(0.0007)** |

**Table 2:** Performance comparison on RNA+ATAC spatial multi-omics benchmark datasets. Values are reported as mean (standard deviation). Best and second-best results are shown in bold and underlined, respectively.

| Methods | ARI | AMI | NMI | FMI | Homo | Purity | Completeness | Jaccard | Dice |
| --- | --- | --- | --- | --- | --- | --- | --- | --- | --- |
| *Mouse Brain* |  |  |  |  |  |  |  |  |  |
| CellCharter | 0.3840(0.0424) | 0.5500(0.0226) | 0.5508(0.0226) | 0.4770(0.0374) | 0.5538(0.0229) | 0.6140(0.0304) | 0.5479(0.0238) | 0.3136(0.0318) | 0.4767(0.0377) |
| SpatialGlue | 0.2946(0.0157) | 0.4295(0.0165) | 0.4305(0.0165) | 0.4090(0.0108) | 0.4249(0.0189) | 0.5514(0.0272) | 0.4364(0.0153) | 0.2566(0.0088) | 0.4084(0.0111) |
| MISO | 0.2000(0.0087) | 0.3160(0.0131) | 0.3173(0.0131) | 0.3538(0.0145) | 0.2846(0.0093) | 0.4520(0.0077) | 0.3591(0.0243) | 0.2099(0.0080) | 0.3469(0.0108) |
| COSMOS | 0.1169(0.0363) | 0.2375(0.0432) | 0.2397(0.0430) | 0.2159(0.0376) | 0.2869(0.0500) | 0.4359(0.0349) | 0.2060(0.0380) | 0.1097(0.0229) | 0.1970(0.0372) |
| PRAGA | 0.2580(0.0542) | 0.3741(0.0703) | 0.3752(0.0702) | 0.3720(0.0474) | 0.3760(0.0689) | 0.5161(0.0456) | 0.3745(0.0716) | 0.2293(0.0361) | 0.3718(0.0474) |
| SMOPCA | 0.2946(0.0101) | 0.4467(0.0051) | 0.4477(0.0051) | 0.4479(0.0120) | 0.4091(0.0058) | 0.5689(0.0108) | 0.4947(0.0106) | 0.2756(0.0078) | 0.4320(0.0097) |
| SMODEL | 0.2255(0.0147) | 0.3857(0.0112) | 0.3868(0.0111) | 0.3504(0.0148) | 0.3790(0.0150) | 0.5076(0.0123) | 0.3951(0.0101) | 0.2121(0.0106) | 0.3499(0.0144) |
| PRESENT | 0.3232(0.0258) | 0.4538(0.0219) | 0.4548(0.0218) | 0.4361(0.0212) | 0.4441(0.0217) | 0.5528(0.0292) | 0.4661(0.0244) | 0.2780(0.0172) | 0.4348(0.0210) |
| SpaMV | 0.2521(0.0206) | 0.3629(0.0171) | 0.3641(0.0171) | 0.3764(0.0101) | 0.3596(0.0215) | 0.5102(0.0237) | 0.3689(0.0150) | 0.2307(0.0081) | 0.3749(0.0107) |
| GatorPrism | **0.5719(0.0571)** | **0.6677(0.0325)** | **0.6684(0.0324)** | **0.6357(0.0486)** | **0.6871(0.0368)** | **0.7919(0.0541)** | **0.6506(0.0306)** | **0.4653(0.0454)** | **0.6351(0.0485)** |
| *Mouse Embryonic-E11.0* |  |  |  |  |  |  |  |  |  |
| CellCharter | 0.3329(0.0412) | 0.4731(0.0402) | 0.4789(0.0397) | 0.4835(0.0360) | 0.5695(0.0456) | 0.7950(0.0278) | 0.4132(0.0355) | 0.2992(0.0316) | 0.4597(0.0378) |
| SpatialGlue | 0.3910(0.0262) | 0.4863(0.0166) | 0.4922(0.0164) | 0.5367(0.0216) | 0.5612(0.0218) | 0.7937(0.0191) | 0.4383(0.0140) | 0.3574(0.0211) | 0.5262(0.0230) |
| MISO | 0.3279(0.0166) | 0.3273(0.0328) | 0.3343(0.0326) | 0.5296(0.0139) | 0.3235(0.0373) | 0.6774(0.0300) | 0.3463(0.0291) | 0.3585(0.0120) | 0.5277(0.0131) |
| COSMOS | 0.2598(0.0465) | 0.3737(0.0278) | 0.3807(0.0275) | 0.4263(0.0373) | 0.4442(0.0324) | 0.6996(0.0285) | 0.3331(0.0244) | 0.2598(0.0303) | 0.4117(0.0373) |
| PRAGA | 0.3690(0.0488) | 0.4759(0.0460) | 0.4819(0.0455) | 0.5183(0.0394) | 0.5524(0.0563) | 0.7806(0.0406) | 0.4274(0.0382) | 0.3395(0.0340) | 0.5060(0.0389) |
| SMOPCA | 0.2786(0.0000) | 0.4069(0.0000) | 0.4137(0.0000) | 0.4536(0.0000) | 0.4646(0.0000) | 0.7213(0.0000) | 0.3728(0.0000) | 0.2894(0.0000) | 0.4489(0.0000) |
| SMODEL | 0.3714(0.0117) | 0.5161(0.0051) | 0.5214(0.0051) | 0.5165(0.0100) | 0.6172(0.0060) | 0.7920(0.0055) | 0.4514(0.0047) | 0.3257(0.0092) | 0.4913(0.0105) |
| PRESENT | 0.4491(0.0501) | 0.5399(0.0232) | 0.5450(0.0229) | 0.5814(0.0411) | 0.6353(0.0254) | 0.8414(0.0159) | 0.4773(0.0218) | 0.3915(0.0408) | 0.5616(0.0429) |
| SpaMV | 0.4116(0.0369) | 0.5322(0.0146) | 0.5374(0.0145) | 0.5507(0.0305) | 0.6259(0.0134) | 0.8140(0.0146) | 0.4709(0.0159) | 0.3634(0.0314) | 0.5324(0.0332) |
| GatorPrism | **0.5219(0.1099)** | **0.5836(0.0846)** | **0.5878(0.0838)** | **0.6404(0.0900)** | **0.6618(0.0932)** | **0.8519(0.0832)** | **0.5286(0.0763)** | **0.4587(0.0803)** | **0.6289(0.0906)** |
| *Mouse Embryonic-E18.5* |  |  |  |  |  |  |  |  |  |
| CellCharter | 0.3701(0.0361) | 0.5705(0.0123) | 0.5781(0.0121) | 0.4605(0.0325) | 0.6289(0.0126) | 0.7176(0.0200) | 0.5349(0.0124) | 0.2875(0.0266) | 0.4460(0.0320) |
| SpatialGlue | 0.4143(0.0757) | 0.5432(0.0187) | 0.5514(0.0184) | 0.4997(0.0668) | 0.5863(0.0199) | 0.6773(0.0126) | 0.5206(0.0191) | 0.3300(0.0589) | 0.4935(0.0679) |
| MISO | 0.2545(0.0288) | 0.3738(0.0102) | 0.3843(0.0104) | 0.4040(0.0275) | 0.3553(0.0127) | 0.5168(0.0143) | 0.4195(0.0216) | 0.2471(0.0190) | 0.3959(0.0249) |
| COSMOS | 0.3321(0.0563) | 0.4966(0.0203) | 0.5055(0.0200) | 0.4260(0.0513) | 0.5533(0.0218) | 0.6424(0.0311) | 0.4653(0.0191) | 0.2586(0.0408) | 0.4094(0.0520) |
| PRAGA | 0.3501(0.0360) | 0.5124(0.0066) | 0.5210(0.0065) | 0.4427(0.0324) | 0.5698(0.0063) | 0.6642(0.0114) | 0.4799(0.0081) | 0.2705(0.0280) | 0.4251(0.0346) |
| SMOPCA | 0.5167(0.0219) | 0.5787(0.0067) | 0.5864(0.0066) | 0.5942(0.0156) | 0.6036(0.0101) | 0.7148(0.0111) | 0.5705(0.0111) | 0.4222(0.0159) | 0.5936(0.0158) |
| SMODEL | 0.3693(0.0199) | 0.5613(0.0087) | 0.5690(0.0086) | 0.4607(0.0180) | 0.6240(0.0093) | 0.7108(0.0140) | 0.5229(0.0081) | 0.2835(0.0152) | 0.4415(0.0185) |
| PRESENT | 0.5465(0.0697) | 0.6174(0.0142) | 0.6245(0.0139) | 0.6171(0.0612) | 0.6462(0.0171) | 0.7341(0.0243) | 0.6044(0.0157) | 0.4473(0.0640) | 0.6156(0.0634) |
| SpaMV | 0.5653(0.0243) | 0.6293(0.0108) | 0.6361(0.0106) | 0.6322(0.0211) | 0.6595(0.0102) | 0.7364(0.0111) | 0.6144(0.0128) | 0.4612(0.0237) | 0.6309(0.0222) |
| GatorPrism | **0.5748(0.0310)** | **0.6516(0.0189)** | **0.6574(0.0186)** | **0.6403(0.0271)** | **0.6943(0.0191)** | **0.7727(0.0296)** | **0.6243(0.0189)** | **0.4621(0.0268)** | **0.6321(0.0274)** |
| *Simulation Dataset (RNA+ATAC)* |  |  |  |  |  |  |  |  |  |
| CellCharter | 0.9230(0.0878) | 0.9487(0.0471) | 0.9491(0.0468) | 0.9466(0.0610) | 0.9565(0.0476) | 0.9757(0.0254) | **0.9433(0.0595)** | 0.9012(0.1116) | 0.9446(0.0642) |
| SpatialGlue | 0.6514(0.1550) | 0.7615(0.0901) | 0.7632(0.0894) | 0.7598(0.1003) | 0.7534(0.1085) | 0.8312(0.0759) | 0.7760(0.0779) | 0.6196(0.1353) | 0.7576(0.1013) |
| MISO | 0.3641(0.0636) | 0.6150(0.0569) | 0.6174(0.0565) | 0.5292(0.0457) | 0.6643(0.0677) | 0.7901(0.0368) | 0.5768(0.0476) | 0.3527(0.0388) | 0.5204(0.0422) |
| COSMOS | 0.8234(0.0373) | 0.8240(0.0349) | 0.8252(0.0346) | 0.8752(0.0254) | 0.8323(0.0427) | 0.9039(0.0339) | 0.8185(0.0289) | 0.7780(0.0392) | 0.8746(0.0252) |
| PRAGA | 0.4819(0.0674) | 0.6554(0.0420) | 0.6575(0.0418) | 0.6168(0.0542) | 0.7212(0.0451) | 0.8577(0.0262) | 0.6044(0.0398) | 0.4293(0.0548) | 0.5988(0.0552) |
| SMOPCA | 0.6537(0.0000) | 0.8349(0.0000) | 0.8360(0.0000) | 0.7500(0.0000) | 0.8853(0.0000) | 0.9313(0.0000) | 0.7919(0.0000) | 0.5873(0.0000) | 0.7400(0.0000) |
| SMODEL | 0.7173(0.0408) | 0.8107(0.0173) | 0.8119(0.0172) | 0.7982(0.0297) | 0.8707(0.0230) | 0.9377(0.0130) | 0.7606(0.0148) | 0.6507(0.0446) | 0.7875(0.0334) |
| PRESENT | 0.4291(0.1705) | 0.6176(0.1497) | 0.6201(0.1488) | 0.5820(0.1292) | 0.6532(0.1538) | 0.7961(0.0954) | 0.5909(0.1452) | 0.4160(0.1405) | 0.5761(0.1297) |
| SpaMV | 0.3122(0.2462) | 0.5524(0.1295) | 0.5555(0.1285) | 0.5340(0.1516) | 0.5568(0.1533) | 0.7496(0.1077) | 0.5563(0.1044) | 0.3753(0.1494) | 0.5310(0.1513) |
| GatorPrism | **0.9555(0.0348)** | **0.9531(0.0268)** | **0.9534(0.0266)** | **0.9688(0.0242)** | **0.9820(0.0313)** | **0.9938(0.0120)** | <u>0.9265(0.0223)</u> | **0.9401(0.0442)** | **0.9687(0.0242)** |

### 4.2.2 GatorPrism improves spatial domain recovery while preserving multi-omic structure on human tonsil slice 2

We evaluated Gator-Prism on human tonsil slice 2 using four annotated tissue domains: tonsillar parenchyma, lymphoid follicle, germinal center, and connective/epithelial tissue. A singleton fifth category was excluded because it lacked a definitive tissue-domain annotation, and all methods were reclustered with *K* = 4. After matching predicted clusters to annotated domains by maximum overlap, GatorPrism achieved the highest ARI of 0.423 (Fig. 2B). This exceeded the strongest competing baseline, SpaMV, by 0.230 ARI points, and outperformed the RNA-only and ADT-only baselines, which achieved ARIs of 0.170 and 0.143, respectively. Other integration methods achieved ARIs between 0.078 and 0.183. GatorPrism more closely recovered the overall tonsil architecture, including the dominant tonsillar parenchyma region and localized lymphoid follicle and germinal center structures. In the UMAP space, GatorPrism showed clearer organization of annotation-associated groups than single-modality and baseline integration methods (Fig. 2C).

We examined whether the integrated representation preserved modality-specific structure. Across repeated subsampling runs, we computed PCCs between pairwise distance matrices from each integrated embedding and the corresponding RNA- and ADT-specific embeddings. GatorPrism achieved median PCCs of 0.574 for RNA and 0.391 for ADT (Fig. 3A). Although SMOPCA and CellCharter showed higher global distance preservation, their *K* = 4 clustering accuracy was lower than that of GatorPrism. Thus, global preservation of single-modality geometry alone was insufficient for accurate domain recovery, whereas GatorPrism balanced modality preservation with domain discrimination. Within the GatorPrism-inferred tissue regions, the fused embedding remained positively correlated with RNA and ADT distance structures across all four clusters (Fig. 3B). After matching the inferred clusters to the annotated tissue domains, RNA and ADT marker heatmaps showed clear region-associated patterns (Fig. 3C,D). Immune-associated RNA and protein markers supported the lymphoid follicle and germinal center regions, whereas stromal and extracellular-matrix-related markers supported the connective/epithelial region. These marker patterns provided multi-omic support for the inferred tissue regions.

**Figure 2:**
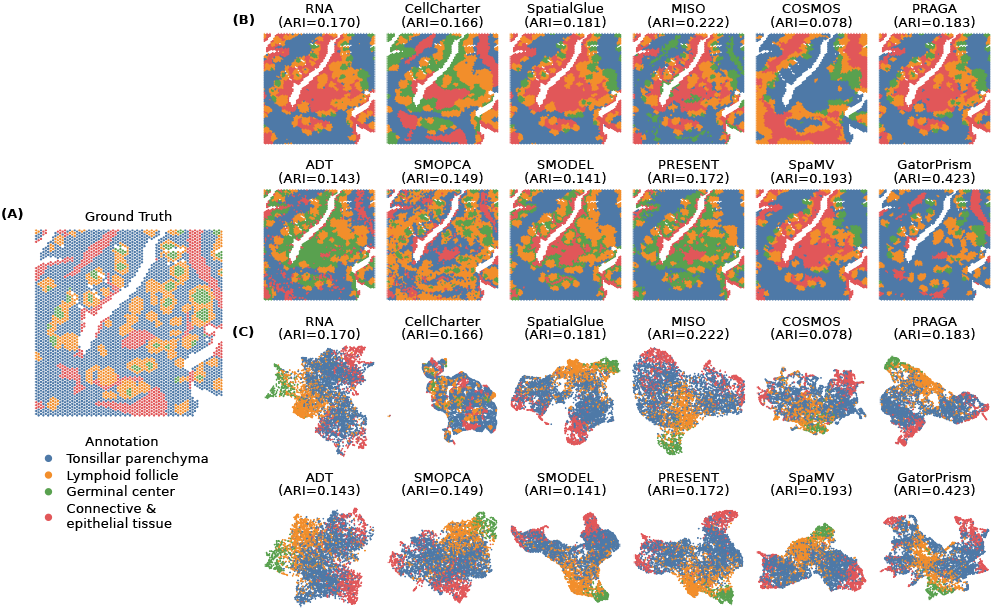
Spatial domain recovery on human tonsil slice 2. (A) Reference tissue annotations. (B) Spatial domain predictions with *K* = 4, with ARI reported in each panel title. (C) UMAP embeddings colored by reference annotation.

**Figure 3:**
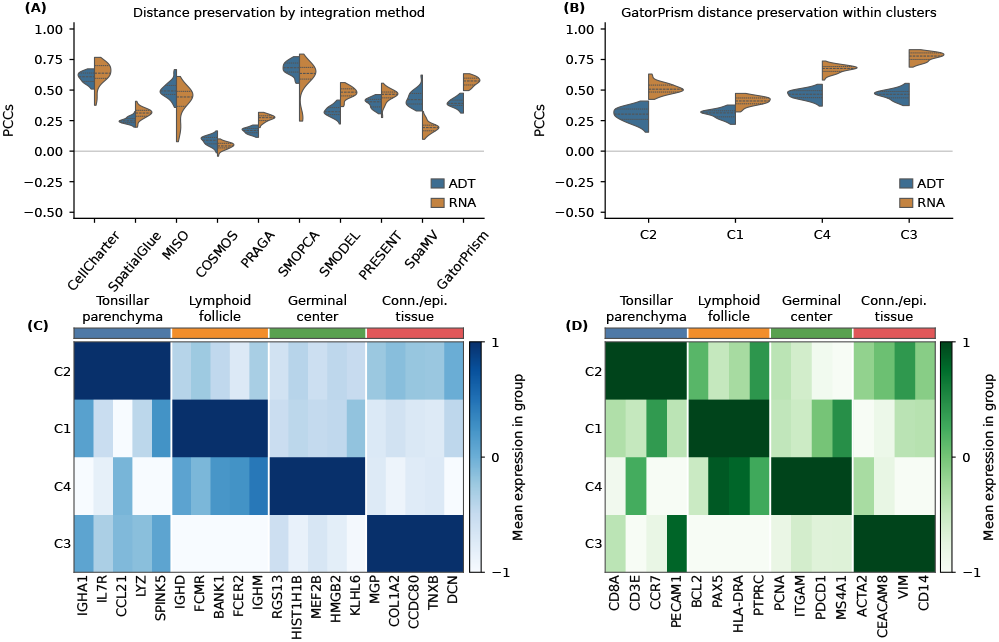
Multi-omic structure preservation and marker patterns in human tonsil slice 2. (A) Global RNA and ADT distance preservation. (B) Cluster-level distance preservation by GatorPrism.. (C,D) RNA and ADT marker expression across matched tissue regions.

### 4.2.3 GatorPrism identifies tissue regions with coherent cross-modality markers and functional programs

Differential ADT and RNA analyses provided biological support for the four matched tissue regions. Representative ADT markers included PAX5 and CXCR5 in lymphoid follicle, PCNA and PDCD1 in germinal center, and ACTA2 and VIM in connective/epithelial tissue (Fig. 4A). Concordant RNA markers included CXCL13 and FCER2, SERPINA9 and RGS13, and MGP and PI16 in the corresponding regions (Fig. 4B). Pairwise analysis further distinguished germinal center from lymphoid follicle, with RGS13, SERPINA9, and MKI67 enriched in the former and BANK1, IGHD, and FCER2 enriched in the latter (Fig. 4C). GO analysis linked germinal-center genes to cell-cycle and chromatin programs, whereas lymphoid-follicle genes were associated with B-cell signaling and immune regulation (Fig. 4D,E). These results support distinct and biologically coherent tissue programs.

**Figure 4:**
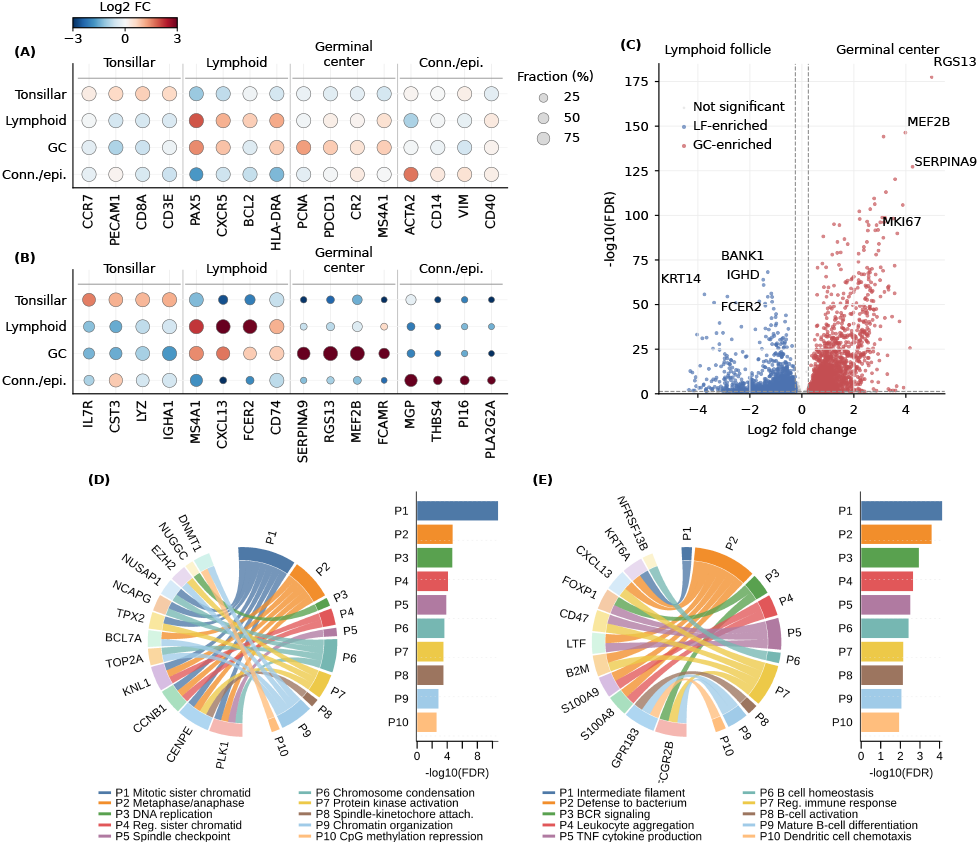
Marker and functional analysis of GatorPrism-inferred regions in human tonsil slice 2. (A,B) ADT and RNA marker dot plots, with color indicating log_2_ fold change and dot size indicating the fraction of expressing spots. (C) Differential expression between the lymphoid follicle and germinal center regions. (D,E) GO enrichment and gene–pathway associations for the two regions.

### 4.2.4 GatorPrism identifies anatomically localized multi-omic routing states in the embryonic mouse brain

We examined whether the learned coalition weights captured anatomical and molecular variation in paired spatial RNA+ATAC data from the E18.5 embryonic mouse brain (Fig. 5). Clustering the GatorPrism representation into 14 domains yielded spatially contiguous regions corresponding to the thalamus (D14; 97%), diencephalon and hindbrain (D4; 95%), and dorsal pallium ventral region (D1; 85%) (Fig. 5A–C). Their routing profiles were distinct: D14 was shared-dominant (0.84), D4 was RNA-private-dominant (0.80), and D1 was ATAC-private-dominant (0.79) (Fig. 5D). Representative molecular patterns supported these routing states (Fig. 5E,F). Ptprd showed concordant RNA and gene-linked ATAC signals in D14, Cntnap2 showed a stronger RNA signal in D4, and Foxo6 showed a stronger accessibility signal in D1. Motif enrichment further distinguished D14 shared-state peaks from D1 ATAC-private peaks (Fig. 5G). In the E11.0–E18.5 comparison, Δ_ATAC_ was defined as the mean ATAC-private routing weight in E18.5 minus that in E11.0, such that positive values indicate greater reliance on ATAC-specific information in E18.5. The difference was positive across all 10 paired training seeds for both the all-spot and neural-only analyses, and in 9 of 10 seeds after adjusting for differences in the composition of shared anatomical regions (Fig. 5H). Shared-anatomy estimates varied between regions and were limited by sparse E11.0 spots, while all-spot and neural-only differences remained positive in balance-loss settings, but the anatomy-adjusted result was less stable.

## 5 Conclusion

We presented GatorPrism, a self-supervised coalition graph mixture-of-experts framework that distinguishes and adaptively integrates cross-omics consensus and modality-specific structure. Across eight RNA+ADT and RNA+ATAC benchmarks, GatorPrism achieved the highest ARI and revealed anatomically localized shared, RNA-private, and ATAC-private states. Future work will extend Gator-Prism to serial-section data and additional modalities.

**Figure 5:**
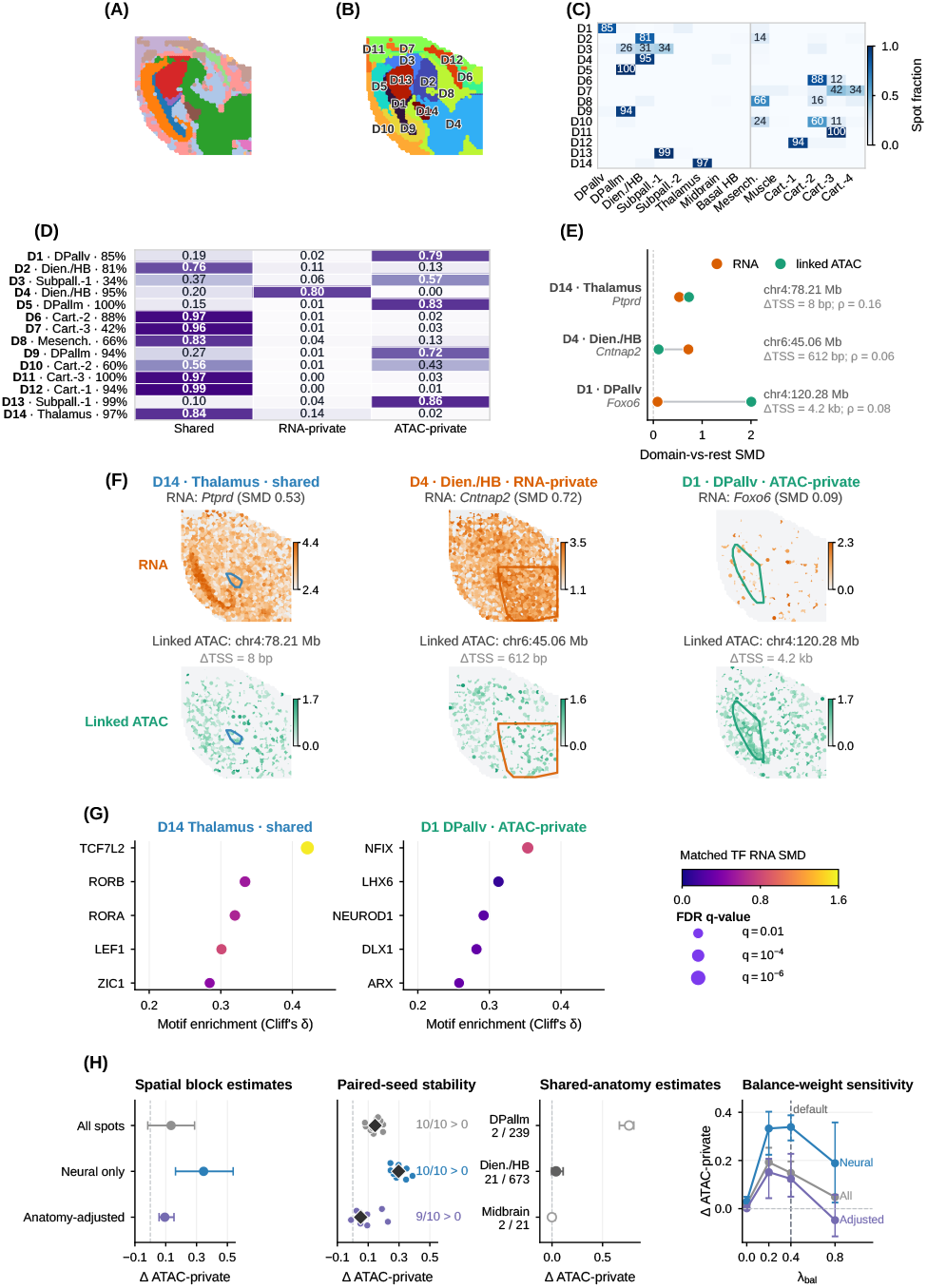
Multi-omic routing in the E18.5 embryonic mouse brain. (A,B) Reference anatomy and inferred domains; (C,D) anatomical overlap and routing profiles; (E,F) representative RNA and linked-ATAC signals; (G) motif enrichment; and (H) E11.0–E18.5 ATAC-private routing comparison and robustness analyses.

## 6 Limitations and Ethical Considerations

Evaluation is limited to co-registered two-modality data, and robustness to additional modalities, strong batch effects, and rare spatial states remains to be established. Human-data applications require appropriate consent, privacy protection, data-use governance, and institutional oversight.

## 7 Generative AI Usage

Generative AI tools were used only for manuscript editing and wording refinement. The authors remain fully responsible for all scientific content.

## A Experimental Setup

### A.1 Benchmark Datasets

Table 3 summarizes the number of spatial units, modality types, feature dimensions, reference domains, platforms, and data sources for all benchmark datasets. The public datasets span different tissues, species, spatial technologies, sample sizes, and modality combinations. The RNA+ADT datasets provide paired transcriptomic and protein-abundance measurements, whereas the RNA+ATAC datasets provide paired transcriptomic and chromatin-accessibility profiles. This benchmark enables a systematic evaluation of GatorPrism across heterogeneous molecular assays and tissue organizations.

**Table 3:**
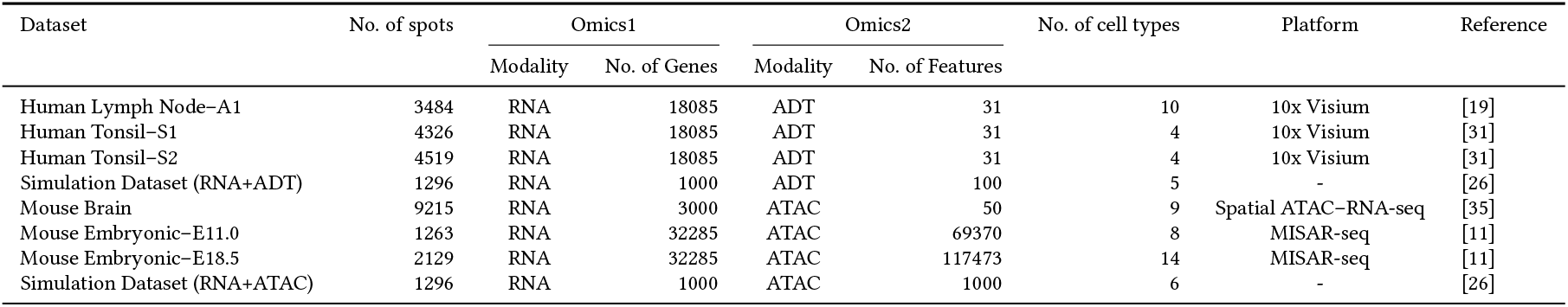
Statistics of the eight spatial multi-omics datasets.

| Dataset | No. of spots | Omics1 |  | Omics2 |  | No. of cell types | Platform | Reference |
| --- | --- | --- | --- | --- | --- | --- | --- | --- |
|  |  | Modality | No. of Genes | Modality | No. of Features |  |  |  |
| Human Lymph Node-A1 | 3484 | RNA | 18085 | ADT | 31 | 10 | 10x Visium | [19] |
| Human Tonsil-S1 | 4326 | RNA | 18085 | ADT | 31 | 4 | 10x Visium | [31] |
| Human Tonsil-S2 | 4519 | RNA | 18085 | ADT | 31 | 4 | 10x Visium | [31] |
| Simulation Dataset (RNA+ADT) | 1296 | RNA | 1000 | ADT | 100 | 5 | - | [26] |
| Mouse Brain | 9215 | RNA | 3000 | ATAC | 50 | 9 | Spatial ATAC-RNA-seq | [35] |
| Mouse Embryonic-E11.0 | 1263 | RNA | 32285 | ATAC | 69370 | 8 | MISAR-seq | [11] |
| Mouse Embryonic-E18.5 | 2129 | RNA | 32285 | ATAC | 117473 | 14 | MISAR-seq | [11] |
| Simulation Dataset (RNA+ATAC) | 1296 | RNA | 1000 | ATAC | 1000 | 6 | - | [26] |

### A.2 Implementation Details

For RNA data, genes detected in fewer than 10 spots were removed, followed by library-size normalization, log transformation, selection of 3,000 highly variable genes, scaling, and PCA. ADT data were processed using centered-log-ratio normalization, scaling, and PCA. For ATAC data, peaks detected in fewer than 10 spots were removed, followed by TF–IDF transformation and LSI, excluding the first LSI component. Symmetrized graphs were constructed with *κ*_*s*_ = 6 spatial neighbors and *κ*_*f*_ = 10 feature neighbors.

Each coalition expert used *L* = 3 graph-attention layers with latent dimension *d* = 64, 4 attention heads, and dropout rate 0.1.

The router and decoders were implemented as two-layer MLPs. We set the Student-*t* parameter and routing temperature to α = τ = 1, and used a contrastive temperature of 0.2. The number of routing prototypes was set to *R* = 5. GatorPrism was trained using AdamW with a learning rate of 10^−3^, weight decay of 10^−4^, and gradient clipping at 5.0. The loss weights were λ_cons_ = 0.3, λ_nbr_ = 0.2, λ_mod_ = 0.5, λ_sm_ = 0.1, λ_bal_ = 0.1, with unit weight for the reconstruction loss.

After training, a *K*-component mclust model was applied to the fused representation Z. For benchmark evaluation, *K* was set to the number of reference domains, but reference labels were not used during representation learning.

GatorPrism was implemented in Python 3.9.23 using PyTorch 2.4.1. Each experiment was repeated with 10 independent random seeds, and results are reported as the mean and sample standard deviation. All experiments were conducted on an NVIDIA RTX 4090 GPU with 24 GB memory.

### A.3 Baseline Methods

To comprehensively evaluate GatorPrism, we selected nine state-of-the-art and representative spatial multi-omics integration methods as baselines, including CellCharter [27], SpatialGlue [19], MISO [6], COSMOS [35], PRAGA [9], SMOPCA [4], SMODEL [15], PRESENT [16], and SpaMV [18].

**CellCharter** identifies spatial cellular niches by combining molecular features with local neighborhood information. It first learns low-dimensional representations and then aggregates features over spatial neighborhoods before clustering cells or spots into spatially coherent niches.

**SpatialGlue** is a graph neural network-based spatial multi-omics integration method with a dual-attention mechanism. It first integrates spatial and feature graphs within each modality and then learns modality-specific attention weights to generate a fused representation for spatial domain identification.

**MISO** is a multimodal spatial omics framework for feature extraction and clustering across diverse data types, including gene expression, protein expression, chromatin accessibility, metabolomics, and histology images. It learns modality-specific embeddings, models cross-modal interactions, and produces integrated features for spatial domain detection.

**COSMOS** integrates paired spatial multi-omics data using two modality-specific graph convolutional encoders, weighted nearest-neighbor fusion, and contrastive learning. It generates an integrated embedding for downstream analyses such as spatial domain segmentation, visualization, and pseudo-spatiotemporal mapping.

**PRAGA** is a prototype-aware graph adaptive aggregation framework for spatial multi-modal omics analysis. It learns dynamic omics-specific graphs to capture latent semantic relations between spots and uses prototype contrastive learning to optimize representations when spot annotations or class-number priors are unavailable.

**SMOPCA** is a spatial multi-omics principal component analysis method for joint dimension reduction. It models multiple omics layers with shared latent factors and incorporates spatial dependencies through covariance structures derived from spatial coordinates.

**SMODEL** is a dual-graph regularized ensemble learning framework for spatial domain identification. It combines anchor concept factorization, element-wise weighted ensemble learning, and graph regularization to learn spatial consensus representations from multiomics data and base clustering results.

**PRESENT** is a contrastive learning-based framework for cross-modality representation and multi-sample integration of spatial omics data. It uses omics-specific encoders, cross-omics alignment, and distribution-aware decoders to integrate spatial transcriptomic, epigenomic, and proteomic modalities.

**SpaMV** is a spatial multi-view representation learning method that explicitly disentangles cross-omics shared information and omics-specific private information. It learns shared and private latent representations for spatial domain clustering, interpretable topic modeling, and omics-specific biomarker discovery.

## B Algorithmic Details

Algorithm 1 summarizes the end-to-end training and inference workflow of GatorPrism. The prototype associations *q*_*ir*_ serve as internal routing variables, whereas the final spatial domains are obtained by applying `mclust` to the fused representation Z.

## C Additional Experimental Analyses

### C.1 Ablation Study

We conducted ablation experiments across eight spatial multi-omics datasets to examine four components of GatorPrism: expert composition, consensus-graph construction, the integration of spatial and molecular edges, and prototype-conditioned routing. The full model was compared with eight variants over ten random seeds using ARI, AMI, NMI, and FMI. Overall, GatorPrism achieved the strongest and most consistent performance across datasets and metrics (Fig. 6). Removing either the joint expert (GatorPrism_*α*_) or the modality-private experts (GatorPrism_*β*_) reduced performance, showing that neither representation alone matches their adaptive integration in the full model. Because GatorPrism_*β*_ contains only one expert, routing and load balancing are no longer applicable; its performance decrease therefore reflects the combined loss of modality-private representations and multi-expert fusion.

**Figure 6:**
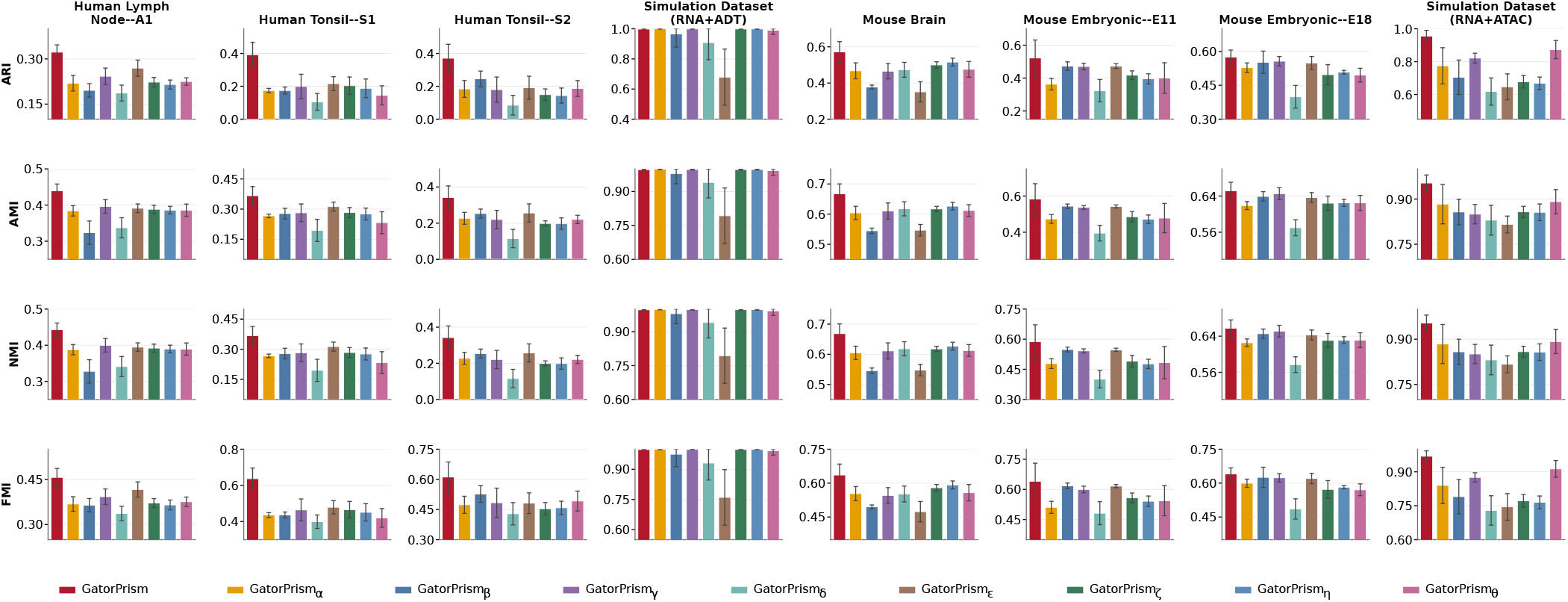
Ablation analysis of GatorPrism across eight spatial multi-omics datasets. Bars show mean performance over ten random seeds, with error bars indicating sample standard deviations. Columns correspond to datasets and rows to ARI, AMI, NMI, and FMI. GatorPrism is the full model; GatorPrism_α_ removes the joint expert; GatorPrism_τ_ retains only the joint expert; GatorPrism_τ_ replaces the intersection-based consensus feature graph with the union graph; GatorPrism_τ_ uses spatial edges only; GatorPrism_τ_ uses feature edges only; GatorPrism_τ_ uses uniform expert weights; GatorPrism_τ_ learns one global gate; and GatorPrism_τ_ uses a spot-specific router conditioned on τ_*i*_ without the prototype-conditioned context τ_*i*_.

#### Algorithm 1

GatorPrism: self-supervised training and inference

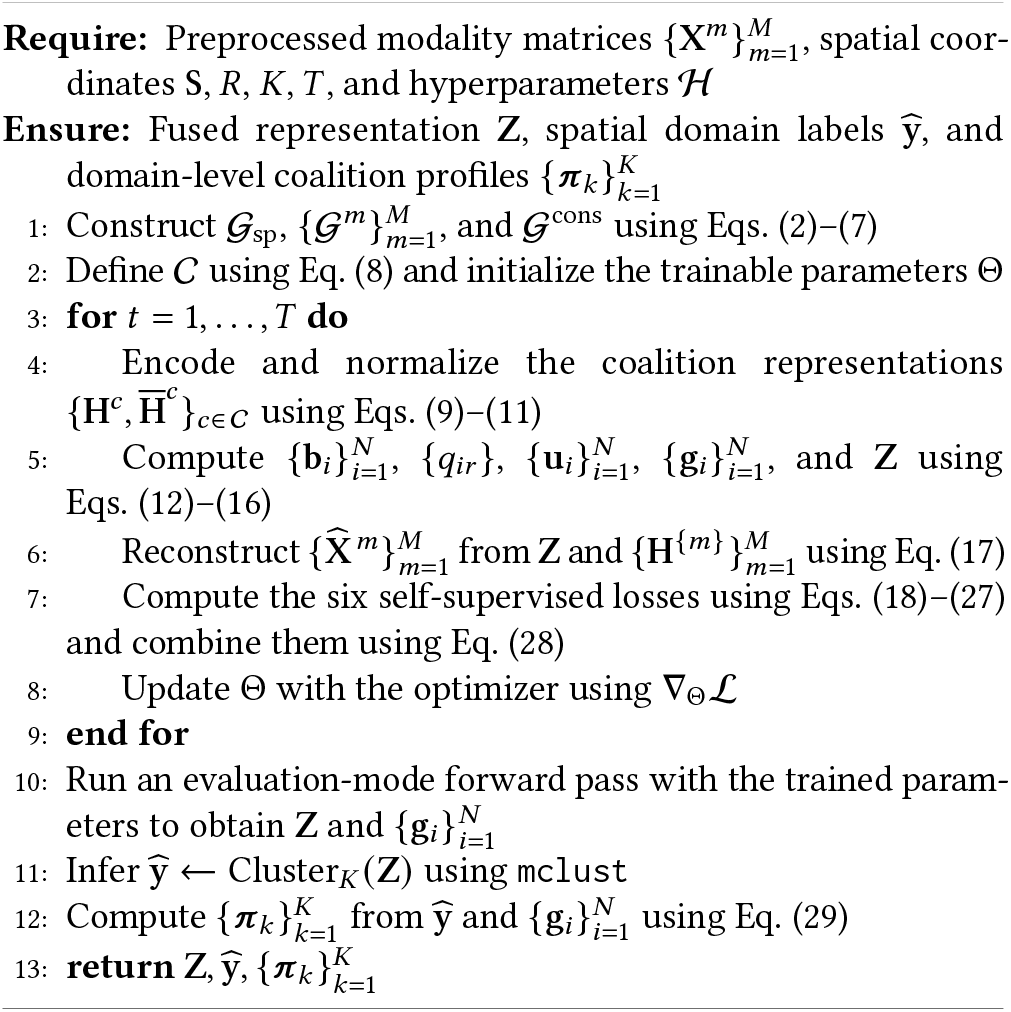

Replacing the intersection-based consensus graph with the union of modality-specific feature graphs (GatorPrism_*γ*_) also reduced performance. This result supports the use of intersection edges as a stricter representation of cross-omics molecular agreement. Similarly, the spatial-only (GatorPrism_*δ*_) and feature-only (GatorPrism_*ϵ*_) variants underperformed the full model, indicating that spatial proximity and molecular similarity provide complementary information for spatial domain identification.

The graph variants further demonstrate the roles of spatial proximity and molecular similarity. GatorPrism_*δ*_ uses spatial edges alone for all experts and therefore removes molecular-neighborhood information, whereas GatorPrism_*ϵ*_ uses feature edges alone and removes explicit spatial proximity. Both variants underperformed the full model, with the decline generally more pronounced for the spatial-only configuration. These results indicate that neither source of neighborhood information is sufficient by itself and that their combination within the joint and modality-specific graphs is important for recovering coherent spatial domains.

Finally, we evaluated three alternative routing strategies while retaining the full expert architecture and training objectives. Uniform routing (GatorPrism_*ζ*_), a single global gate (GatorPrism_*η*_), and spot-specific routing without the prototype-conditioned context (GatorPrism_*θ*_) all performed worse than the full model. In particular, GatorPrism_*θ*_ provides the most direct test of prototype conditioning because it preserves spot-specific routing but excludes *u*_*i*_ . Its consistent performance gap relative to GatorPrism suggests that prototype-derived context improves adaptive expert routing across heterogeneous spatial multi-omics datasets.

### C.2 Hyperparameter Sensitivity Analysis

We evaluated the robustness of GatorPrism across eight spatial multi-omics datasets by varying five key hyperparameters: spatial neighbors, feature neighbors, GAT attention heads, dropout rate, and MoE gate temperature. For each setting, performance was averaged over five random seeds using ARI, AMI, NMI, and FMI. GatorPrism maintained stable performance across hyperparameter ranges (Fig. 7). Graph-construction parameters had the largest effect: moderate spatial and feature neighborhoods generally yielded stronger performance, whereas overly dense neighborhoods tended to reduce accuracy, likely due to over-smoothing or cross-domain mixing. Dropout also showed a clear regularization effect, with moderate dropout improving average performance across datasets. In contrast, GAT attention heads and MoE gate temperature had relatively smaller effects, indicating that GatorPrism does not depend on a narrowly tuned attention or routing configuration. Overall, these results demonstrate that GatorPrism remains robust across hyperparameter settings and supports its applicability to diverse spatial multi-omics datasets.

**Figure 7:**
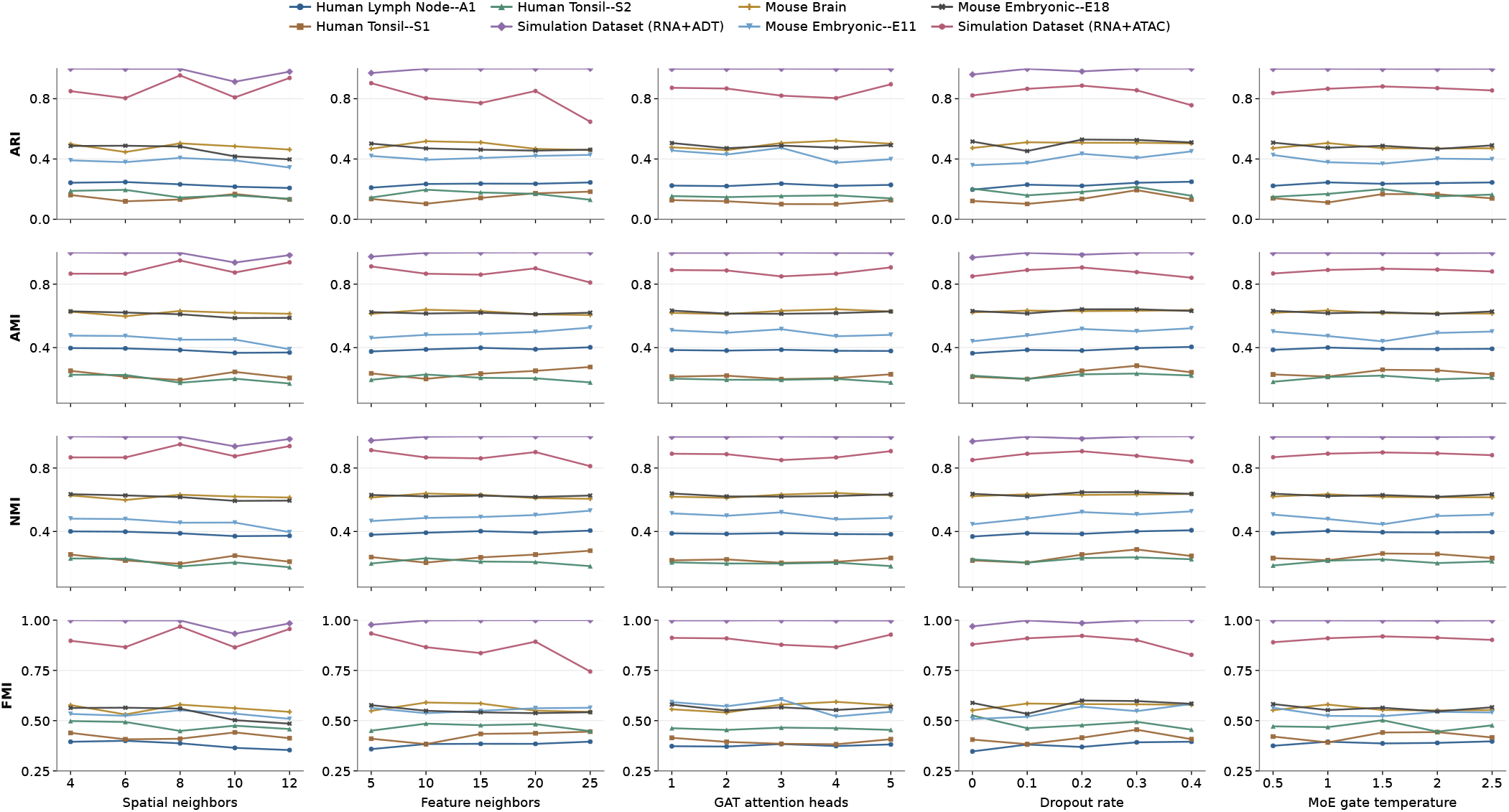
Hyperparameter sensitivity of GatorPrism across eight spatial multi-omics datasets.

### C.3 Accuracy–Efficiency Analysis

We evaluated the accuracy–efficiency trade-off of GatorPrism on two RNA+ATAC brain benchmarks, including Mouse Brain and Mouse Embryonic–E18.5, by comparing ARI against training time and GPU memory usage, as shown in Fig. 8. On the Mouse Brain dataset, GatorPrism achieved the highest ARI of 0.578, exceeding the strongest competing baseline, CellCharter, by 0.190 ARI points. It required 28.86 seconds for training, which was comparable to Spatial-Glue and MISO, and was faster than COSMOS, PRAGA, SMOPCA, SMODEL, PRESENT, and SpaMV. Its GPU memory usage was 3.12 GB, remaining within the same practical range as most competing methods while requiring less memory than SMOPCA and SMODEL. On Mouse Embryonic–E18.5, GatorPrism again achieved the highest ARI of 0.596, outperforming SpaMV, PRESENT, and SMOPCA, which achieved ARIs of 0.563, 0.543, and 0.519, respectively. GatorPrism also maintained efficient runtime and memory usage on this dataset, requiring 18.3 seconds for training and 0.49 GB of GPU memory. This made it faster than most competing methods, including SpatialGlue, MISO, COSMOS, PRAGA, SMODEL, PRESENT, and SpaMV, while using less GPU memory than MISO, COSMOS, PRAGA, SMODEL, PRESENT, and SpaMV. These results suggest that GatorPrism improves spatial domain identification accuracy on RNA+ATAC brain benchmarks without incurring excessive computational cost.

**Figure 8:**
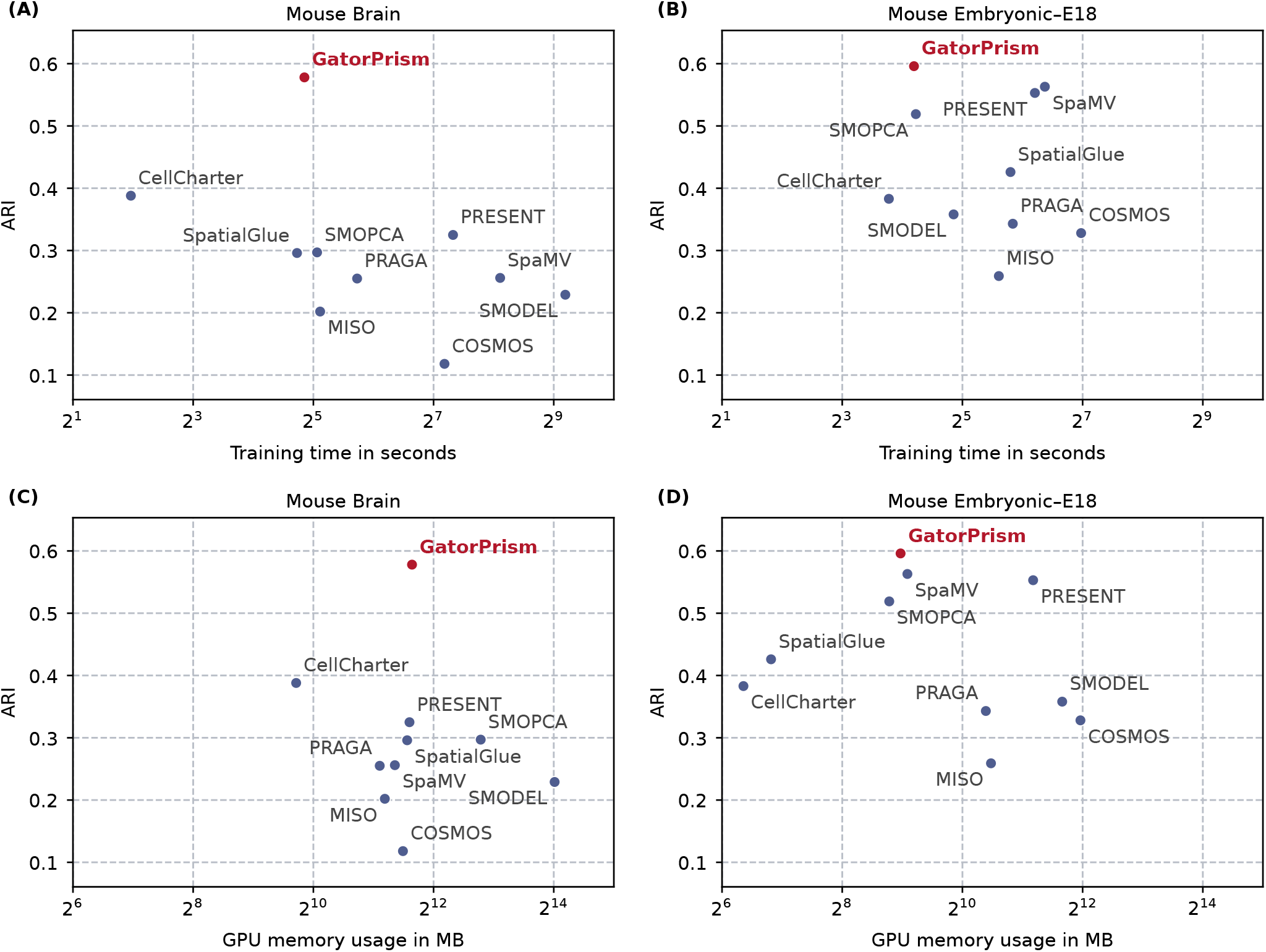
Accuracy–efficiency trade-off on RNA+ATAC mouse brain benchmarks. ARI is compared against training time and GPU memory usage on Mouse Brain and Mouse Embryonic–E18.5. (A,B) ARI versus training time. (C,D) ARI versus GPU memory usage.

